# Influence of basal media composition on barrier fidelity within human pluripotent stem cell-derived blood-brain barrier models

**DOI:** 10.1101/2021.03.01.433282

**Authors:** Emma H. Neal, Ketaki A. Katdare, Yajuan Shi, Nicholas A. Marinelli, Kameron A. Hagerla, Ethan S. Lippmann

**Affiliations:** Department of Chemical and Biomolecular Engineering, Vanderbilt University, Nashville, TN, USA; Vanderbilt Brain Institute, Vanderbilt University, Nashville, TN, USA; Chemical and Physical Biology Program, Vanderbilt University, Nashville, TN, USA; Department of Biomedical Engineering, Vanderbilt University, Nashville, TN, USA; Department of Neurology, Vanderbilt University Medical Center, Nashville, TN, USA

**Keywords:** Blood-brain barrier, *in vitro* model, brain microvascular endothelial cell, human induced pluripotent stem cell

## Abstract

It is increasingly recognized that brain microvascular endothelial cells (BMECs), the principle component of the blood-brain barrier (BBB), are highly sensitive to soluble cues from both the bloodstream and the brain. This concept extends *in vitro*, where the extracellular milieu can also influence BBB properties in cultured cells. However, the extent to which baseline culture conditions can affect BBB properties *in vitro* remains unclear, which has implications for model variability and reproducibility, as well as downstream assessments of molecular transport and disease phenotypes. Here, we explore this concept by examining BBB properties within human induced pluripotent stem cell (iPSC)-derived BMEC-like cells cultured under serum-free conditions in different basal media with fully defined compositions. We demonstrate notable differences in both passive and active BBB properties as a function of basal media composition. Further, RNA sequencing and phosphoproteome analyses revealed alterations to various signaling pathways in response to basal media differences. Overall, our results demonstrate that baseline culture conditions can have a profound influence on the performance of *in vitro* BBB models, and these effects should be considered when designing experiments that utilize such models for basic research and preclinical assays.

## Background

Brain microvascular endothelial cells (BMECs), which comprise the functional component of the blood-brain barrier (BBB), respond dynamically to a variety of soluble cues. A plethora of *in vitro* BBB models, constructed with cells derived from primary bovine, porcine, rodent, and human sources, have demonstrated alterations to BMEC barrier functions upon non-contact co-culture with other cell types or treatment with proteins and small molecules (reviewed extensively in (Helms *et al*. 2016)). More recently, a series of *in vivo* studies have characterized the dynamic behavior of BMECs in response to complex changes in plasma composition (Chen *et al*. 2020; Yousef *et al*. 2019). Yet, when reflecting on how to properly study such *in vivo* phenomena in more controlled *in vitro* BBB models, a persistent limitation stands out: *in vitro* models predominantly use undefined culture conditions, particularly animal serum. This issue limits the standardization of model performances and can lead to unexpected variability. Such variability can have significant implications on interpretations of preclinical assays, for example predictions of small molecule and biological drug transport across the BBB. As such, our goal herein was to better understand how the composition of cell culture media can influence passive and active barrier properties within *in vitro* BBB models.

To study this issue, we utilized a well-characterized model system that employs endothelial-like cells with BBB properties (termed BMEC-like cells) differentiated from human pluripotent stem cells (hPSCs). As first described in 2012 (Lippmann *et al*. 2012), hPSC-derived BMEC-like cells express canonical endothelial and BBB markers, form a passive barrier as measured by transendothelial electrical resistance (TEER), express a cohort of active efflux transport proteins, and exhibit representative permeability to well-characterized small molecules that correlates well with *in vivo* uptake in rodents. Retinoic acid (RA) treatment was subsequently shown to upregulate VE-cadherin expression, as well as significantly increase TEER and multidrug resistance protein (MRP) activity (Lippmann *et al*. 2014a). Since the publication of these initial studies, as well as others that validated the BBB phenotype (Katt *et al*. 2016; DeStefano *et al*. 2017; Vatine *et al*. 2019; Delsing *et al*. 2018; Linville *et al*. 2019; Lim *et al*. 2017; Canfield *et al*. 2017), hPSC-derived BMEC-like cells have been incorporated into increasingly complex neurovascular models ranging from 2D Transwell setups to 3D microfluidic chips that incorporate neurons, astrocytes, and pericytes from either primary or iPSC sources (Wang *et al*. 2016; Canfield *et al*. 2019; Appelt-Menzel *et al*. 2017; Vatine *et al*. 2019; Park *et al*. 2019; Brown *et al*. 2020; Motallebnejad *et al*. 2019). Such models have proven useful for analyzing diseases that afflict the neurovascular unit (Lim *et al*. 2017; Vatine *et al*. 2017; Blanchard *et al*. 2020). They have also gained traction for use in preclinical drug screening applications (Di Marco *et al*. 2020; Ohshima *et al*. 2019; Roux *et al*. 2019). Yet, like animal BBB models, the use of hPSC-derived BMEC-like cells is not standardized, and differences in cellular function and phenotype have been noted across the literature, which could reflect a combination of cell sourcing, technique, and culture conditions (Lippmann *et al*. 2020). Therefore, further efforts are warranted to improve understanding of how various external factors influence hPSC-derived BBB performance *in vitro*, and lessons learned from these studies could be potentially translated to other *in vitro* model platforms.

Ultimately, we decided to focus this study on basal media composition. We recently described a series of improvements to the canonical hPSC-to-BMEC differentiation procedure that removed many undefined components in the cell culture media, particularly knockout serum replacement and platelet-poor serum (Hollmann *et al*. 2017; Neal *et al*. 2019). Overall, these changes vastly improved the consistency of differentiation and certain properties in the BMEC-like cells (for example, responsiveness to co-culture with astrocytes). However, the basal medium used in the later stages of differentiation and after purification of the BMEC-like cells (human endothelial serum-free medium, or hESFM) has a proprietary composition. Thus, despite the removal of serum from the differentiation procedure, we still had a limited understanding of the exact baseline cues the BMEC-like cells were receiving and how they were responding to these cues. To this end, we examined replacement of hESFM with either Neurobasal Medium (NB) or Dulbecco’s Modified Eagle Medium/Nutrient Mixture F12 (DMEM/F12, referred to hereafter as DMEM). In contrast to hESFM, the full compositions of NB and DMEM are publicly available through their manufacturer. Both media contain a similar profile of amino acids, vitamins, nutrients, and inorganic salts, but at varying compositions, thus providing a useful testbed for assessing how seemingly innocuous cues might influence the BBB phenotype *in vitro*.

Through a variety of standard *in vitro* BBB assays, we show that BMEC-like cells exhibit different passive and active barrier properties when cultured in NB versus DMEM basal media. RNA sequencing revealed differential regulation of a number of genes, including ones that encode molecular transporters. A KEGG pathway analysis highlighted potential key differences in signaling activity, which were further validated by assessing the phosphorylation status of several kinases. Overall, our results help benchmark the plasticity and responsiveness of hPSC-derived BMEC-like cells to their extracellular environment and provide a new avenue for studying how soluble cues influence BBB properties using completely defined media compositions. More generally, these findings may aid in the standardization and interpretation of BBB properties in models constructed not only from hPSCs, but also from primary and immortalized cells.

## Methods

### hPSC maintenance

hPSCs were maintained as previously described (Hollmann *et al*. 2017; Neal *et al*. 2019). CC3 iPSCs (Kumar *et al*. 2014), CD10 iPSCs (Tidball *et al*. 2016), and CDH5-2A-eGFP knock-in reporter H9 hESCs (Bao *et al*. 2017) were maintained in E8 medium (Chen *et al*. 2011), prepared as previously detailed (Hollmann *et al*. 2017). hPSCs were cultured on tissue culture plates coated with growth-factor reduced Matrigel (Corning #354230). hPSCs were passaged with Versene (Fisher Scientific #15-040-066) upon reaching 60-80% confluency. All hPSCs utilized in this study were from cryopreserved stocks previously determined to be karyotypically normal, and hPSCs were not used for experiments after passage 40.

### hPSC differentiation to BMEC-like cells

hPSCs were differentiated to BMEC-like cells as previously described, with minor modifications (Neal *et al*. 2019). hPSCs were washed once with PBS (Fisher Scientific #14-190-144), dissociated with Accutase (Fisher Scientific #A1110501) for 3 minutes, and collected by centrifugation. hPSCs were then resuspended in E8 medium containing 10 μM Y27632 (Tocris #1254) and seeded onto Matrigel-coated 6-well plates at a density of 15,800 cells/cm^2^. The following day, the cells were switched to E6 medium (Lippmann *et al*. 2014b) to initiate differentiation. Media was changed every day for 4 days. On day 4, the cells were switched to EC medium, which consisted of a basal medium supplemented with 200× B27 (Fisher Scientific #17-504-044), 10 μM all-trans retinoic acid (RA; Sigma-Aldrich #R2625), and 20 ng/mL basic fibroblast growth factor (bFGF; PeproTech #100-18b). The basal medium was human endothelial serum-free medium (hESFM; Thermo Fisher Scientific #11111044), neurobasal medium (NB; Thermo Fisher Scientific #21103049), or DMEM/F12 with Glutamax (DMEM; Thermo Fisher Scientific #10565018). Cells were then left to incubate for 48 hours in EC medium without a media exchange. On day 6, resultant BMEC-like cells were purified and assayed as described below. For the CDH5-2A-eGFP line, fluorescent images were acquired at day 6 on live cells using a Leica DMi8 microscope with an environmental chamber.

### Purification of hPSC-derived BMEC-like cells

On day 6 of differentiation, hPSC-derived BMEC-like cells were purified similar to previous descriptions (Lippmann *et al*. 2012; Lippmann *et al*. 2014a; Hollmann *et al*. 2017; Neal *et al*. 2019). Briefly, cells were washed once with DPBS, incubated with Accutase until a single cell suspension was achieved (approximately 20-45 min), and collected via centrifugation. Cells were resuspended in EC medium and plated onto Transwell filters (Fisher Scientific #07-200-161), cell culture plates (Corning), or 35 mm dishes with number 1.5 coverslips (MatTek Corporation #P35G1.514C) coated with 400 μg/mL collagen IV (Sigma-Aldrich #C5533) and 100 μg/mL fibronectin (Sigma-Aldrich #F1141). In all experiments, BMEC-like cells were purified using the same basal medium as during differentiation (e.g. if cells were differentiated in EC medium made from DMEM, they were subsequently subcultured into EC medium made from DMEM). Media was changed approximately 24 hours after subculture to EC medium lacking bFGF and RA. In certain experiments, BMEC-like cells were changed to different basal media containing 200× B27 at specified time points – these experiments are noted in the text. With the exception of these select experiments, medium was not changed again after the removal of bFGF and RA for the duration of culture.

### TEER measurements

After purification of BMEC-like cells onto Transwell filters, TEER was measured approximately every 24 hours using a World Precision Instruments EVOM2 voltohmmeter with STX2 chopstick electrodes.

### Fluorescein permeability measurements

Three Transwell filters plated with BMEC-like cells, as well as one Transwell filter coated with collagen IV and fibronectin but containing no cells, were used to test permeability for each basal medium condition. All permeability assays were conducted 24 hours after removal of bFGF and RA from EC medium, referred to as day 1 of subculture. One hour prior to commencement of the assay, BMEC-like cells on Transwells received a full medium change in both apical and basolateral compartments. At the start of the assay, medium was fully aspirated from the apical portion of all Transwell filters and replaced with EC medium containing 10 μM sodium fluorescein (Sigma-Aldrich #F6377). 200 μL of medium was immediately removed from the basolateral chamber of each filter, transferred to a 96-well plate, and replaced with 200 μL of fresh medium containing no fluorescein. This process was repeated every 30 minutes for a total of five time points with cells incubated at 37°C between medium removal and replacements. At the conclusion of the assay, the fluorescence of the collected samples was measured using a Tecan Infinite M1000Pro microplate reader, and effective permeability (Pe) values were calculated as previously described (Stebbins *et al*. 2016).

### Transporter activity assays

Transporter activity was measured on day 1 of subculture. For each transporter studied, BMEC-like cells were incubated with fluorescent transporter substrate with or without transporter inhibitors. For inhibitor conditions, BMEC-like cells in well plates were treated with 10 μM cyclosporin A (CsA; Fisher Scientific #11-011-00), a p-glycoprotein inhibitor, or 10 μM MK571 (Sigma-Aldrich #M7571), an MRP inhibitor, for 1 hour on a rocking platform at room temperature. After 1 hour, medium was aspirated from all wells and replaced with medium containing 10 μM rhodamine 123 (R123; Thermo Fisher Scientific #R302), a p-glycoprotein substrate, or 10 μM 2’,7’-dichlorodihydrofluorescein diacetate (H2DCFDA; Thermo Fisher Scientific #D399), an MRP family substrate, with or without their respective inhibitors. After a 1 hour incubation at 37°C, BMEC-like cells were washed three times with PBS. Three wells of BMEC-like cells per condition were then lysed in PBS containing 5% Triton X-100 (Sigma-Aldrich #T8787), and one well per condition was fixed in ice cold 100% methanol. The fluorescence of the collected cell lysates was measured using a Tecan Infinite M1000Pro microplate reader. Cells fixed in methanol were incubated with 4’,6-Diamidino-2-pheny-lindoldihydrochloride (DAPI; Thermo Fisher Scientific #D1306) for 10 minutes and subsequently imaged at 6 locations per condition using a Leica DMi8 microscope. Nuclei at each location were counted using Fiji (Schindelin *et al*. 2012) and used to determine cell density per well of each condition. Measured fluorescence values were subsequently normalized on a per cell basis using these counts.

### Immunocytochemistry

Cells were washed twice with PBS before fixation in 4% paraformaldehyde (PFA; Fisher Scientific #AAJ19943K2) for 20 minutes or ice-cold 100% methanol for 10 minutes. Following fixation, cells were washed with PBS three times for five minutes per wash. Cells fixed in 4% PFA were blocked in PBS containing 5% donkey serum (Sigma-Aldrich #D9663) and 0.3% Triton X-100 (Sigma-Aldrich), collectively referred to as PBS-DT, for a minimum of 1 hour. Cells fixed in methanol were blocked in PBS with 5% donkey serum but lacking 0.3% Triton X-100, referred to as PBSD, for a minimum of 1 hour prior to primary antibody addition. Primary antibodies were diluted in either PBS-DT or PBSD, depending on the buffer used to block the cells, to the appropriate dilution (Table S1), and incubated overnight at 4°C.

Following overnight incubation, cells were rinsed once with DPBS and washed for an additional five times, 5 minutes per wash. Secondary antibodies (Table S2) were diluted 1:200 into PBS-DT or PBSD and incubated for 1-2 hours at room temperature. Nuclei were labeled in a 10 minute incubation using DAPI or Hoechst 33342 trihydrochloride trihydrate (Thermo Fisher Scientific #H1399) diluted in PBS-T or PBS. Finally, cells were rinsed once and washed four times, 5 minutes per wash, and imaged using a Leica DMi8 microscope.

### RNA sequencing and pathway analysis

On day 1 of subculture, BMEC-like cells that had been plated on 6-well cell culture plates coated with collagen IV and fibronectin were washed once with PBS and collected via scraping and centrifugation. Resultant cell pellets were resuspended in 500 μL of Trizol (Thermo Fisher Scientific #15596026) and stored at −80°C. After collection of 3 biological replicates per media condition, RNA from each sample was isolated using a Direct-zol Miniprep kit (Zymo Research #R2051), following the manufacturer’s instructions with inclusion of a DNase treatment step. RNA samples were submitted to the Vanderbilt Technologies for Advanced Genomics (VANTAGE) facility for sequencing using an Illumina NovaSeq6000. Raw sequencing reads were obtained for the 6 paired-end samples and run through a bulk RNA-Seq pipeline governed by the Snakemake (5.2.4) workflow management system (Köster and Rahmann 2018). Quality control was monitored with FastQC v0.11.8 before and after quality and adapter trimming, as performed by Trim Galore v0.5.0. Trimmed paired-end sequences were then aligned to the human genome (GRC38) with STAR 2.60c, utilizing Gencode genomic feature annotations (v26). Once quantitated, the feature counts were used for downstream analysis. The average mapped reads count was 41M reads (85% of total reads). Gene counts and sample metadata were used for differential gene expression in DESeq2 (Love *et al*. 2014). Principal Component Analysis and hierarchical clustering revealed that one of the DMEM samples was an outlier, so it was removed from downstream analysis. For each testing pair, we analyzed differential gene expression results quantitatively and visually. For the up- and down-regulated groups of top genes, we also performed functional gene enrichment analysis for Gene Ontology and KEGG pathways.

### Phospho-proteome analysis

The relative phosphorylation levels of various kinases were measured using a Proteome Profiler Array human phospho-kinase array kit (R&D Systems #ARY003C) according to the manufacturer’s instructions. Briefly, on day 1 of subculture, BMEC-like cells were washed once with DPBS, incubated with Accutase, and collected via centrifugation at 1000 rpm for 4 minutes. Cells were resuspended in 1 mL of DPBS, and cell density was determined using a Countess II. Cells were again centrifuged at 1000 rpm for 4 minutes and resuspended in Lysis Buffer 6 at a concentration of 10^7^ cells/mL. The resultant cell suspensions were incubated on ice for 30 minutes, followed by centrifugation at 14,000×*g* for 5 minutes. Supernatants were transferred to clean microfuge tubes, and protein content was determined using a Pierce BCA protein assay kit (Thermo Fisher Scientific #23225). Lysates were stored at −80°C until the day of the assay. On the day of the assay, all kit components were prepared as directed by the manufacturer’s instructions, and 400 μg of protein per condition was diluted in Array Buffer 1 for incubation with kit membranes. Chemiluminescence for all membranes was detected using a Li-Cor Biosciences Fc imaging system at exposure times of 30 seconds, 2 minutes, and 10 minutes. Data collected from the 2 minute exposure were selected for presentation as this time maximized signal intensity while minimizing background signal.

### Western blot analysis

On day 1 of subculture, BMEC-like cells were washed with 1 mL of DPBS, collected by scraping and transferred into a 15 mL conical in 1 mL of DPBS, and pelleted at 1000 rpm for 4 minutes. Cell pellets were resuspended in RIPA buffer (Sigma-Aldrich # R0278) with 1% v/v protease inhibitor cocktail (Sigma-Aldrich #P8340) and 1% v/v phosphatase inhibitor cocktail 3 (Sigma-Aldrich #P0044) and transferred to clean microfuge tubes. Lysates were incubated on ice for 30 minutes. After this incubation, lysates were centrifuged for 15 minutes at 12000×*g* at 4°C, and cleared supernatants were transferred to clean microfuge tubes. Protein isolates not immediately used were stored at −20°C.

Protein concentration for all samples was determined using a Pierce BCA protein assay. Each sample was prepared for western blotting by diluting 20 μg of protein with 1 × Laemmli buffer (Bio-Rad #1610747) and ultrapure water (Invitrogen #10977015) to a final volume of 20 μL per sample. Prepared samples were heated at 95°C for 5 minutes, placed on ice for an additional 5 minutes, and briefly centrifuged. Samples, as well as Precision Plus Protein Dual Color standard (Bio-Rad #1610374), were loaded into 4-20% Criterion TGX precast midi protein gels (Bio-Rad #5671094). Gels were run at 80V until all protein samples had fully entered the gel. Voltage was then increased to 160V until desired degree of protein separation was obtained. Protein was transferred to a nitrocellulose membrane (Thermo Fisher Scientific # IB23001) using an iBlot 2 gel transfer device (Thermo Fisher Scientific #IB21001), and membranes were blocked in Intercept (TBS) blocking buffer (Li-Cor Biosciences # 92760001) for a minimum of 30 minutes at room temperature on an orbital shaker. Following blocking, membranes were incubated with primary antibodies (Table 3) diluted in Intercept (TBS) blocking buffer with 0.05% Tween-20 (Sigma-Aldrich) overnight at 4°C on an orbital shaker.

Membranes were rinsed once with 1× TBS-T, prepared by diluting 10× TBS (Fisher Scientific # MT46012CM) with distilled water and adding 0.05% Tween-20 (Sigma-Aldrich #P9416), and washed three times, 5 minutes per wash, in TBS-T. The secondary antibody (IRDye 800CW goat anti-rabbit; Li-Cor #926-32211) was diluted in TBS-T at 1:15,000 and incubated with membranes for 1-2 hours at room temperature on an orbital shaker. Membranes were rinsed once and washed three times with TBS-T for 5 minutes per wash. Membranes were imaged using a Li-Cor Odyssey CLx Imager, and band intensity was determined using Image Studio Lite software.

### Ethical statement and study design

Institutional ethical approval was not required for this study. Pre-registration and randomization were not required for this study. No blinding was performed for experimental analyses. No sample calculations were performed. This study was exploratory with no inclusion or exclusion criteria. For all experiments, a biological replicate refers to an independent iPSC differentiation. When iPSC-derived BMEC-like cells are purified into multiple Transwell filters or wells for analysis, each of the filters or wells is considered a technical replicate.

### Statistical analyses

GraphPad Prism software was used for statistical analyses of wet lab experiments. As specified in each figure legend where statistical analyses were performed, the student’s unpaired t-test, one-way ANOVA, and two-way ANOVA were used to assess statistical significance. Normality of the data was not assessed. The ROUT test revealed no outliers.

## Results

### Marker expression in different basal media

hPSCs were differentiated for 4 days in E6 medium and 2 days in EC medium containing varying basal media of hESFM, DMEM, or NB (Figure 1A). The standard additives to EC medium (RA, B27 supplement, bFGF) were unchanged from our previous protocol (Neal *et al*. 2019). In cells differentiated from the H9 CDH5-2A-eGFP hESC line, expression of eGFP was similar at day 6 regardless of basal media, and colonies of cells harboring green fluorescence signal at cell-cell borders could be identified in each condition (Figure 1B); these results agree with previous observations (Lippmann *et al*. 2020). For further validation, we differentiated CC3 and CD10 iPSCs in either DMEM or NB, followed by subculture and immunostaining. In all cases, the BMEC-like cells maintained a VE-cadherin+ endothelial signature with smooth claudin-5+ and occludin+ tight junctions, as well as expression of GLUT-1 (Figure 1C). Thus, on a qualitative level, the use of different basal media did not impact the differentiation procedure or the gross identity of the purified BMEC-like cells.

**Figure 1.**
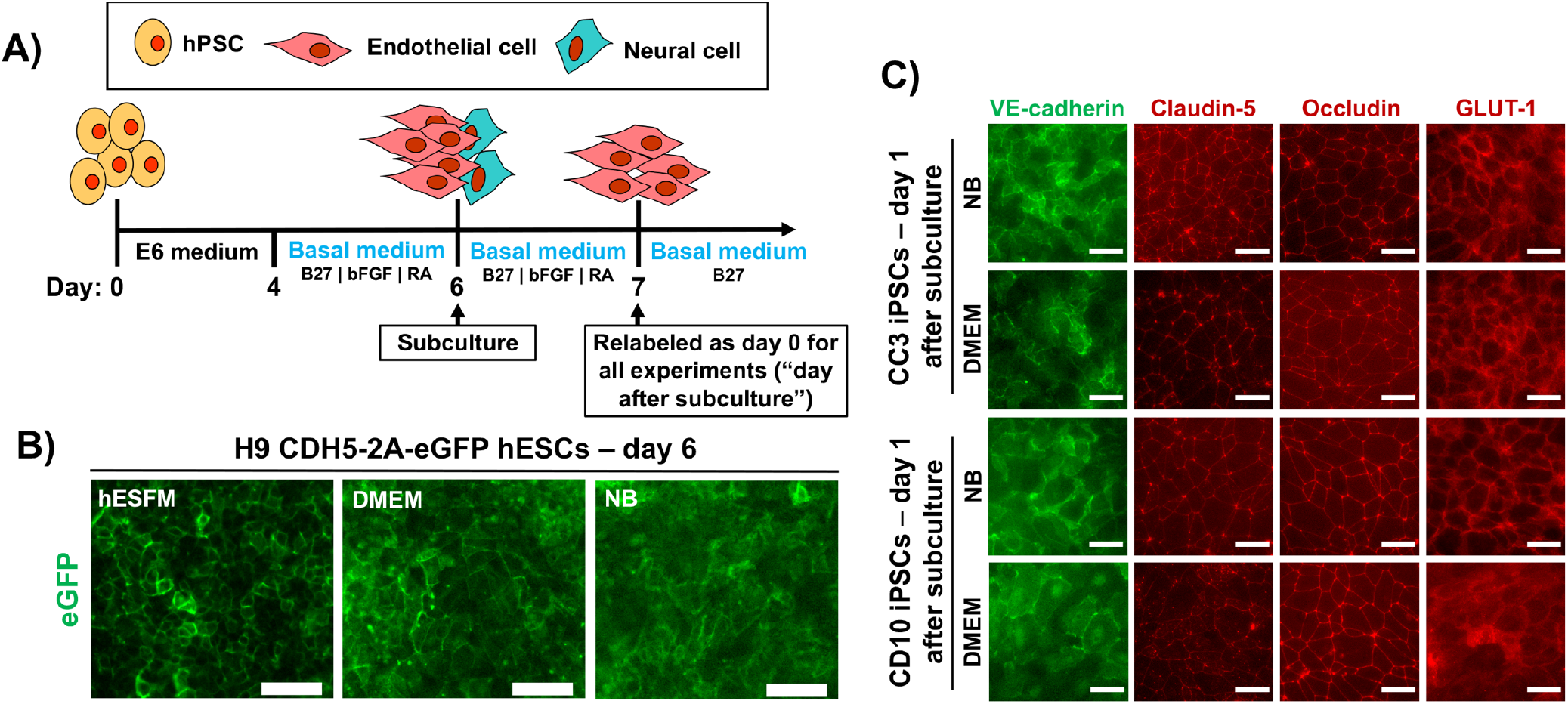
Study design and assessment of characteristic endothelial and BBB markers. (A) Timeline for differentiation of hPSCs into BMEC-like cells. Basal medium is the only altered variable between all experiments. The day of subculture is noted, as well as the notation used for experiments that occur after subculture. (B) eGFP fluorescence as a function of basal media in the H9 CDH5-2A-eGFP hESC line at day 6 of differentiation. (C) Expression of VE-cadherin, claudin-5, occludin, and GLUT-1 in purified BMEC-like cells at day 1 after subculture. Representative images are shown in BMEC-like cells derived from two different iPSC lines (CC3 and CD10). In all images, scale bars are 50 μm.

### Passive barrier function in response to different basal media

We next assessed passive barrier function in the BMEC-like cells using measurements of TEER and small molecule permeability. Whereas BMEC-like cells differentiated from CC3 iPSCs and subcultured in NB exhibited TEER in line with maximum values previously observed in hESFM (7,000-8,000 Ωxcm^2^) (Neal *et al*. 2019), TEER values in BMEC-like cells differentiated and subcultured in DMEM were consistently and significantly lower (2,000-4,000 Ωxcm^2^) (Figure 2A). These relative differences were confirmed in BMEC-like cells derived from CD10 iPSCs (Figure 2B). To clarify whether these results were simply suboptimal differentiations (e.g. if differentiation in DMEM produces unhealthy cells because a critical factor is missing from the medium), we characterized the dynamic responsiveness to media composition. We first differentiated and subcultured BMEC-like cells in DMEM medium, then switched to NB medium after 1 day (Figure 2C) or 7 days (Figure 2D). In both cases, the switch to NB medium yielded a statistically significant increase in TEER, showcasing that diminished passive barrier function in DMEM was reversible and not the result of an intrinsic defect acquired during the differentiation process. Similarly, differentiation and subculture in NB medium, followed by a switch to DMEM medium, also yielded a statistically significant decrease in TEER, though the effect was less pronounced (Figure 2E). Despite these dramatic differences in TEER, sodium fluorescein permeability was not significantly different between BMECs in either DMEM or NB medium (Figure 2F), but this result was unsurprising given that *in vitro* permeability differences are not observed in Transwell assays when TEER exceeds 500-1,000 Ωxcm^2^ (Mantle *et al*. 2016). Overall, our data suggest that basal media can influence the relative strength of paracellular barrier function.

**Figure 2.**
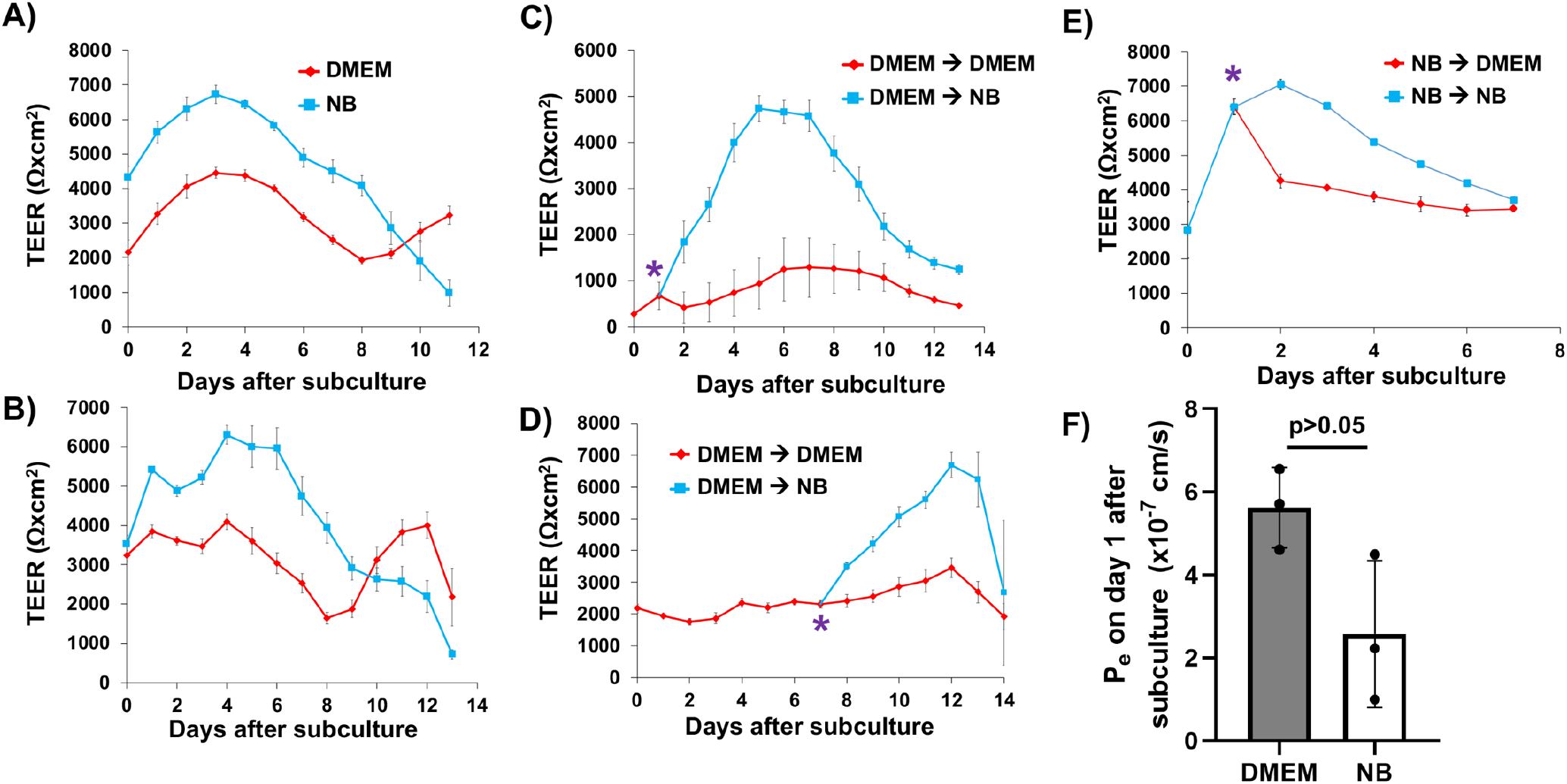
Passive permeability in iPSC-derived BMEC-like cells as a function of basal media. All TEER plots represent a single biological replicate. Each time point is the result of a technical N=9 as cell are purified onto triplicate filters for all conditions tested, and each filter was measured in 3 locations per time point. All data are represented as mean ± standard deviation for these collective measurements. Trends were then confirmed for the specified number of biological replicates. (A) Representative TEER in CC3-derived BMEC-like cells differentiated in NB versus DMEM media. These trends were confirmed across more than 10 biological replicates. (B) Representative TEER in CD10-derived BMEC-like cells differentiated in NB versus DMEM media. These trends were confirmed across 4 biological replicates. (C-E) Representative TEER plots in CC3-derived BMEC-like cells in response to the dynamic media changes indicated in each panel. The asterisk indicates the time point at which the media was changed. The trends in each plot were confirmed across 3 biological replicates. (F) Effective permeability coefficients for sodium fluorescein in CC3-derived BMEC-like cells differentiated in NB versus DMEM media. Data are presented as mean ± standard deviation from triplicate Transwell filters for each condition (a single biological replicate). A student’s unpaired t-test was used to evaluate the difference in fluorescein Pe between the media conditions. Trends were confirmed in two additional biological replicates.

### Basal media influence on transcriptome and transporter regulation

Given the rather striking differences in passive BBB function between BMEC-like cells differentiated using NB and DMEM, we employed RNA sequencing to identify differentially regulated genes to gain insight into these phenotypic differences. RNA was collected from BMEC-like cells differentiated in either DMEM or NB, sequenced and analyzed for differential expression using DESeq2 (Love *et al*. 2014). Our analysis identified 19,086 genes with a nonzero total read count within these samples, and differential expression was found to cluster strongly as a function of basal media condition (Figure 3A shows Principle Component Analysis and Table S4 shows the full list). Using an adjusted *p*<0.05, we identified 2,793 genes as being upregulated (15%) and 3,069 genes as being downregulated (17%) under DMEM conditions as compared to NB conditions. Of these upregulated genes, 490 genes were upregulated at least 2-fold under DMEM conditions and 97 genes were at least 5-fold upregulated under DMEM conditions (Figure 3B). 680 genes were downregulated a minimum of 2-fold under DMEM conditions, and 192 genes were downregulated a minimum of 5-fold under DMEM conditions (Figure 3B). A number of differentially expressed genes encoded for transporters, including efflux transporters and solute carriers; we cross-referenced this list with a single-cell RNA-sequencing database of brain endothelial cells (Vanlandewijck *et al*. 2018) to identify predicted *in vivo* BBB transporters, and indeed several of these BBB transporters were differentially regulated *in vitro* by media conditions, including *ABCB1* (p-glycoprotein), *ABCC4* (MRP4), *SLC1A1, SLC2A1* (GLUT1), *SLC6A6, SLC7A5* (LAT1), and *SLC39A8* (Figure 3C). We followed up on these results using well-established accumulation assays to measure efflux transporter activity, specifically probing p-glycoprotein activity using rhodamine 123 (substrate) and cyclosporin A (inhibitor) and multidrug resistance proteins using 2’,7’-dichlorodihydrofluorescein diacetate (H2DCFDA, substrate) and MK571 (inhibitor). We determined that rhodamine 123 accumulation was significantly increased by cyclosporin A treatment in cells cultured in DMEM but not NB conditions (Figure 3D). We also found that H2DCFDA accumulation was significantly increased by MK571 treatment in cells cultured under either DMEM or NB conditions, but the increase was more substantial under DMEM conditions (Figure 3E). Collectively, these results show that basal media can influence the expression of various genes, including transporters, as well as the activity of these transporters.

**Figure 3.**
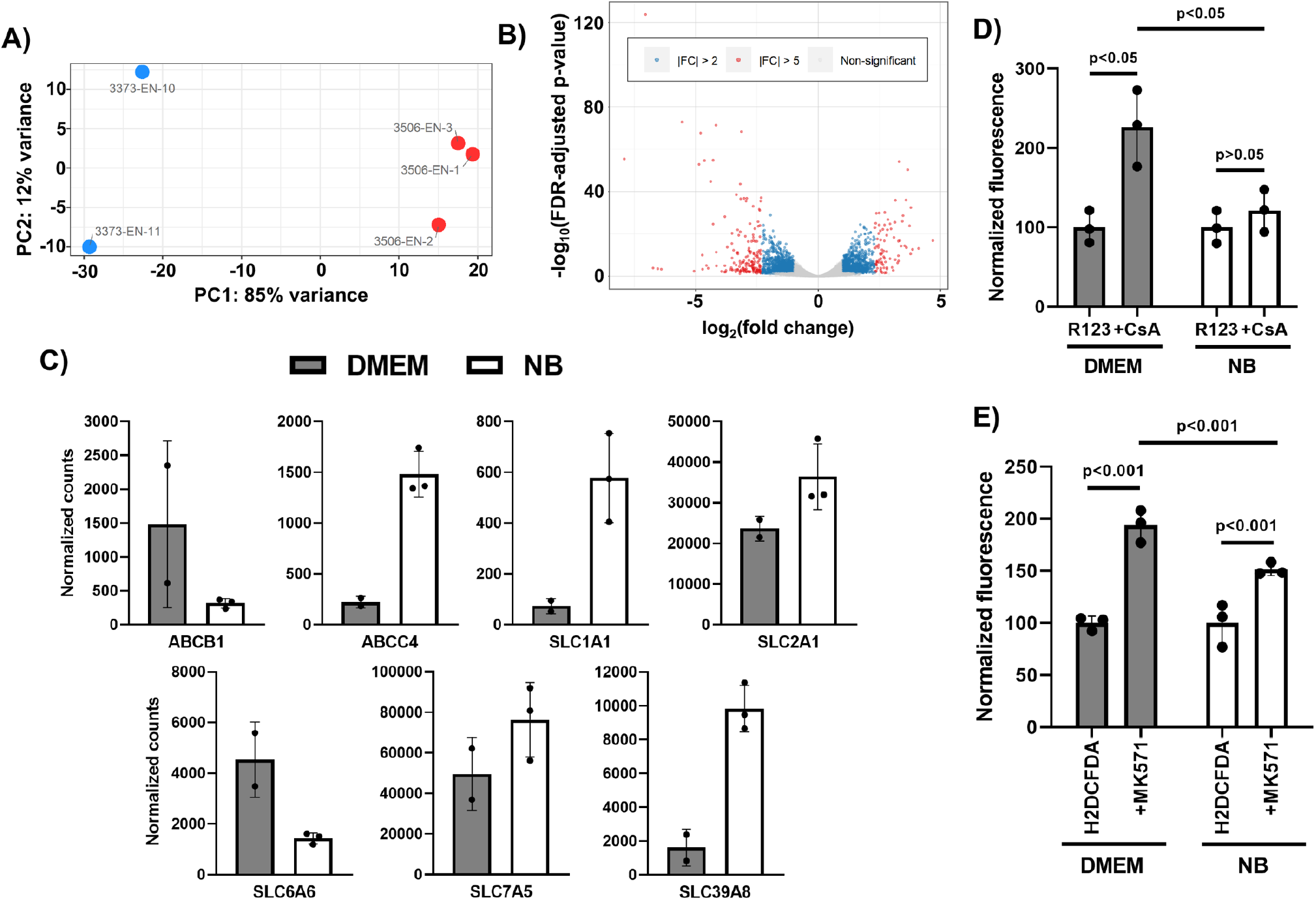
Basal media alters the transcriptome and transporter activity in iPSC-derived BMEC-like cells. (A) Principle Component Analysis of the DMEM and NB samples. The red dots are NB samples and the blue dots are DMEM samples. (B) Volcano plot highlighting up- and downregulated genes in DMEM versus NB. Blue dots indicate genes with a fold-change greater than 2 and red dots indicate genes with a fold-change greater than 5. (C) Normalized counts for select transporters. Data represent mean ± standard deviation from biological replicates. Raw and normalized counts for all genes are found in Table S4. (D-E) Substrate accumulation assays for p-glycoprotein (panel D) and MRPs (panel E). Fluorescence was measured on a per cell basis in three separate wells (technical triplicate), and within each media condition, values are presented as mean ± standard deviation normalized to cells treated with substrate only. A two-way ANOVA was used to evaluate statistical significance upon inhibitor treatment, as well as the degree of increase upon inhibitor treatment between the two media conditions. The trends between media conditions and statistically significant differences were validated in an additional biological replicate.

### Basal media influence on intracellular signaling

We further employed a pathway analysis to understand cellular alterations in response to basal media, which revealed seven downregulated KEGG pathways and 39 upregulated KEGG pathways under DMEM conditions relative to NB conditions (*q*<0.1) (Figure 4A shows select list and Table S5 shows full list). The nature of some of these pathways suggested potential changes to intracellular signaling in response to basal media. To query this possibility, we performed two semi-quantitative, antibody-based assays: a commercially available phospho-kinase array and western blotting with phospho-specific antibodies. In both cases, we detected changes in the phosphoproteome. The phospho-kinase array showed numerous changes to the phosphorylation status of various kinases, including ERK1/2, GSK-3, and β-catenin, which were more heavily phosphorylated under NB conditions (a select portion of the array is shown in Figure 4B and the entire array is shown in Figure S1). We further explored specific changes to ECM-receptor and focal adhesion signaling pathways by western blotting for phosphorylation status of ERK1/2 and FAK, which showed differential regulation (relevant bands are shown in Figure 4C and full blots are provided in Figure S2). Collectively, these results highlight that basal media can influence signaling pathway activation in BMEC-like cells, which in turn may alter BBB properties.

**Figure 4.**
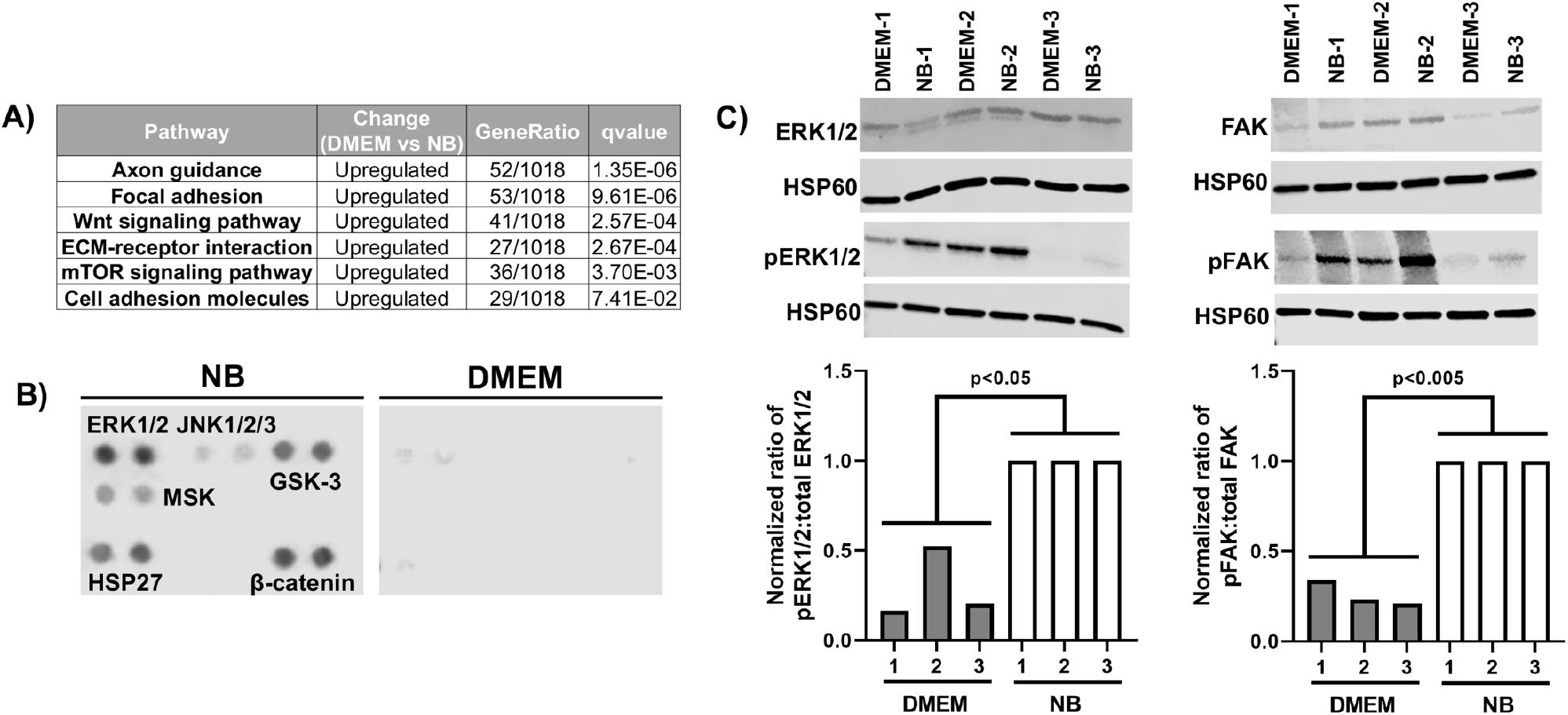
Basal media alters signaling pathways in iPSC-derived BMEC-like cells. (A) Select differentially altered KEGG pathways. The full list of pathways is found in Table S5. (B) Analysis of protein phosphorylation using a phosphoproteome array. Based on how the assay is run, this analysis represents a biological N=1, with technical duplicates, for each media condition. A relevant subset of the array is shown here, and the full array is found in Figure S1. (C) Assessment of total and phospho-specific expression of ERK1/2 and FAK by western blot. Samples 1, 2, and 3 represent biological replicates where the corresponding DMEM and NB samples were prepared from the same iPSC differentiation. Each blot was first normalized to HSP60. Ratios of phosphorylated to total protein were then normalized to the NB condition within each pair of samples (NB-1 to DMEM-1, NB-2 to DMEM-2, and NB-3 to DMEM-3). Each biological replicate is presented as an individual bar on the graph. The entire cohort was analyzed for statistical significance using a one-way ANOVA. Uncropped images of all western blots are found in Figure S2.

## Discussion

In this study, we explored how basal media composition influences BBB properties in iPSC-derived BMEC-like cells. We determined that differences in basal media, which can include baseline concentrations of salts, amino acids, and vitamins, alter both the passive and active BBB phenotype. RNA sequencing and protein analyses revealed many differentially expressed genes and phosphorylated kinases. Our results benchmark the performance of iPSC-derived BMEC-like cells in different media and will serve as a resource in this area moving forward. Our results also suggest that media composition may be an important consideration when interpreting experimental results in other BBB model formats.

These findings are likely important for a variety of *in vitro* applications ranging from drug screening to disease modeling. For instance, in drug screening applications, proper assessment of molecular transport (which can be impacted by transporter expression and activity levels) is critical. Our data show that, although TEER was above the minimal threshold necessary to prevent paracellular leakage, different basal media yielded notable changes to transporter expression in the iPSC-derived BMEC-like cells. Thus, small molecule permeability measurements could vary widely depending on media composition and its effects on molecular transporters, including both efflux transporters and solute carriers (which can influence drug influx (Abdullahi *et al*. 2017)), without any direct relevance to BBB function *in vivo*. When human *in vivo* data for the BMEC transcriptome ultimately become available, it will be useful to benchmark the transcriptomes of the *in vitro* models against these results, which will assist in selecting the best media for *in vitro* studies as well as aid in designing new media that promotes more *in vivo*-like characteristics within the *in vitro* models.

One interesting outcome from the pathway analyses in this study was the possible identification of differential Wnt signaling between the media conditions. Canonical Wnt/β-catenin signaling is important for CNS angiogenesis and the development of BBB properties (Daneman *et al*. 2009; Stenman *et al*. 2008; Liebner *et al*. 2008). In the canonical Wnt signaling pathway, β-catenin is constitutively phosphorylated on the N-terminus by GSK-3β, which targets the protein for ubiquitination and degradation; inhibition of GSK-3β and/or stabilization of β-catenin by other means allows the protein to translocate to the nucleus and act as a transcription factor (Verheyen and Gottardi 2010). The KEGG pathway analysis suggests that the NB condition yields a lower level of Wnt signaling relative to the DMEM condition. In the phosphoproteome array, β-catenin phosphorylation was elevated in the NB condition, indeed suggesting degradation and downregulation of Wnt/β-catenin signaling. Phosphorylation of GSK-3 was also elevated in the NB condition at either serine-9 (GSK-3α) and serine-21 (GSK-3β); the antibody in the array did not discriminate between these two isoform sites. These modifications are associated with decreased active site availability and inhibition of GSK-3-mediated phosphorylation of substrates (Beurel *et al*. 2015), which potentially disagrees with the β-catenin phosphorylation status. However, it is possible that phosphorylation of GSK-3α, but not GSK-3β, is being detected by this antibody. Other phosphorylation sites can also enhance the activity of GSK isoforms, for example tyrosine-216 and tyrosine-279 (Beurel *et al*. 2015), and antibodies to detect these modifications were not present in the array. Moreover, other signaling pathways can feed into the activity of GSK, and many different pathways were more active under NB conditions. Thus, it is possible that the basal media alter intracellular signaling and subsequently barrier functions through the Wnt/β-catenin pathway, but more work is needed to fully interrogate these findings, potentially through more nuanced molecular biology techniques that utilize phospho-specific antibodies and small molecule-mediated pathway manipulation. Future work may also involve assessment of which BBB properties are specifically impacted through this signaling pathway versus others, as well as whether these effects are mediated through canonical or non-canonical Wnt pathways (the latter being independent of β-catenin).

Given that the two media compositions tested in this study primarily differ by amino acid and vitamin content, we initially believed that the differences in BBB properties between the basal media conditions could also be the result of changes to metabolic pathways within the cells as a function of nutrient availability. The KEGG analysis does suggest this notion is possible, given that the mTOR signaling pathway was differentially regulated. Other differentially regulated pathways that seemed relevant to BBB and endothelial cell function (besides the previously discussed Wnt/β-catenin signaling) included ECM-receptor interactions, focal adhesions, and axon guidance. The behavior of iPSC-derived BMEC-like cells has previously been shown to be strongly dependent upon matrix composition (Al-Ahmad *et al*. 2019; Katt *et al*. 2018), which aligns with our observed connections between BBB phenotype and differential regulation of ECM-receptor and focal adhesion signaling pathways. The identification of differential axon guidance pathway signaling is perhaps more surprising, but there is previously described interplay between developing vasculature and neural cells (Paredes *et al*. 2018), and axon guidance signaling has previously been identified using molecular profiling of mouse retinal microvascular cells (Jeong *et al*. 2017). These observations do not discount that differences in signaling pathway activity could be due to nutrient availability and metabolic feedback, but more nuanced techniques like metabolic tracing may be necessary to explicitly establish these links.

Overall, we have demonstrated that basal media composition can influence both passive and active barrier properties in hPSC-derived BMEC-like cells. Moving forward, these outcomes and technological developments may assist a number of future studies. For example, we have characterized BBB function and transcriptional identity under two fully defined media compositions. These newer, serum-free systems can be utilized for explicit cause-and-effect experiments to study how any potential factor (nutrients, metabolites, hormones, etc.) may influence the BBB phenotype. The knowledge for how basal media influence BBB properties might also be used to improve the function of primary cell-based models, which historically exhibit a diminished BBB phenotype after isolation and *in vitro* culture. Finally, the ability to utilize fully defined, serum-free culture conditions could improve *in vitro* drug transport predictions by mimicking differential concentrations of soluble cues under healthy and diseased conditions. These considerations could extend towards modeling natural biological processes that endogenously fluctuate, for example, circadian rhythms that impact BBB transporter functions (Zhang *et al*. 2018; Cuddapah *et al*. 2019) or hormonal changes associated with stress that could impact drug permeation through the BBB (Brown *et al*. 2020).

## Acknowledgments

Funding was provided by a Ben Barres Early Career Acceleration Award from the Chan Zuckerberg Initiative (grant 2018-191850 to ESL), National Institutes of Health grants R21 NS106510 (to ESL), R01 NS110665 (to ESL), R61 NS112445 (to ESL) and P30 DK020593 (pilot and feasibility award to ESL), and National Science Foundation grant 1846860 (to ESL). EHN was supported by a Graduate Research Fellowship from the National Science Foundation (DGE-1445197). NAM was supported by the Integrated Training in Engineering and Diabetes Program (T32 DK101003). Support for RNA sequencing was provided by the Vanderbilt VANTAGE core facility, which is supported in part by a Clinical and Translational Science Award (5UL1 RR024975), the Vanderbilt Ingram Cancer Center (P30 CA68485), the Vanderbilt Vision Center (P30 EY08126), a CTSA award from the National Center for Advancing Translational Sciences (UL1 TR002243), and the National Center for Research Resources (G20 RR030956). Support for transcriptome analyses was provided by Creative Data Solutions, a Vanderbilt shared resource. The authors would like to thank Lawrence Hsu and Dr. Jean-Philippe Cartailler for helpful discussions on RNA sequencing experiments. The authors declare that they have no competing interests.

## Authors’ contributions

EHN and ESL conceived the study. EHN, KAK, NAM, YS, KAH, and NAM performed the experiments. EHN and ESL wrote the manuscript. All authors read and approved the final manuscript.

## Availability of data and materials

The datasets generated during the current study are available at ArrayExpress under accession ID E-MTAB-8802.

## Notes

https://www.ebi.ac.uk/arrayexpress/experiments/E-MTAB-8802/

### Competing Interest Statement

The authors have declared no competing interest.

## References

Abdullahi W., Davis T. P., Ronaldson P. T. (2017) Functional Expression of P-glycoprotein and Organic Anion Transporting Polypeptides at the Blood-Brain Barrier: Understanding Transport Mechanisms for Improved CNS Drug Delivery? AAPS J. 19, 931–939.

Al-Ahmad A. J., Patel R., Palecek S. P., Shusta E. V. (2019) Hyaluronan impairs the barrier integrity of brain microvascular endothelial cells through a CD44-dependent pathway. J. Cereb. Blood Flow Metab. Off. J. Int. Soc. Cereb. Blood Flow Metab. 39, 1759–1775.

Appelt-Menzel A., Cubukova A., Günther K., Edenhofer F., Piontek J., Krause G., Stüber T., Walles H., Neuhaus W., Metzger M. (2017) Establishment of a Human Blood-Brain Barrier Co-culture Model Mimicking the Neurovascular Unit Using Induced Pluri- and Multipotent Stem Cells. Stem Cell Rep. 8, 894–906.

Bao X., Bhute V. J., Han T., Qian T., Lian X., Palecek S. P. (2017) Human pluripotent stem cell-derived epicardial progenitors can differentiate to endocardial-like endothelial cells. Bioeng. Transl. Med. 2, 191–201.

Beurel E., Grieco S. F., Jope R. S. (2015) Glycogen synthase kinase-3 (GSK3): regulation, actions, and diseases. Pharmacol. Ther. 148, 114–131.

Blanchard J. W., Bula M., Davila-Velderrain J., Akay L. A., Zhu L., Frank A., Victor M. B., et al. (2020) Reconstruction of the human blood–brain barrier in vitro reveals a pathogenic mechanism of APOE4 in pericytes. Nat. Med. 26, 952–963.

Brown J. A., Faley S. L., Shi Y., Hillgren K. M., Sawada G. A., Baker T. K., Wikswo J. P., Lippmann E. S. (2020) Advances in blood-brain barrier modeling in microphysiological systems highlight critical differences in opioid transport due to cortisol exposure. Fluids Barriers CNS 17, 38.

Canfield S. G., Stebbins M. J., Faubion M. G., Gastfriend B. D., Palecek S. P., Shusta E. V. (2019) An isogenic neurovascular unit model comprised of human induced pluripotent stem cell-derived brain microvascular endothelial cells, pericytes, astrocytes, and neurons. Fluids Barriers CNS 16, 25.

Canfield S. G., Stebbins M. J., Morales B. S., Asai S. W., Vatine G. D., Svendsen C. N., Palecek S. P., Shusta E. V. (2017) An Isogenic Blood-Brain Barrier Model Comprising Brain Endothelial Cells, Astrocytes and Neurons Derived from Human Induced Pluripotent Stem Cells. J. Neurochem. 140, 874–888.

Chen G., Gulbranson D. R., Hou Z., Bolin J. M., Ruotti V., Probasco M. D., Smuga-Otto K., et al. (2011) Chemically defined conditions for human iPSC derivation and culture. Nat. Methods 8, 424–429.

Chen M. B., Yang A. C., Yousef H., Lee D., Chen W., Schaum N., Lehallier B., Quake S. R., Wyss-Coray T. (2020) Brain Endothelial Cells Are Exquisite Sensors of Age-Related Circulatory Cues. Cell Rep. 30, 4418–4432.e4.

Cuddapah V. A., Zhang S. L., Sehgal A. (2019) Regulation of the blood-brain barrier by circadian rhythms and sleep. Trends Neurosci. 42, 500–510.

Daneman R., Agalliu D., Zhou L., Kuhnert F., Kuo C. J., Barres B. A. (2009) Wnt/beta-catenin signaling is required for CNS, but not non-CNS, angiogenesis. Proc Natl Acad Sci U A 106, 641–646.

Delsing L., Dönnes P., Sánchez J., Clausen M., Voulgaris D., Falk A., Herland A., et al. (2018) Barrier Properties and Transcriptome Expression in Human iPSC-Derived Models of the Blood–Brain Barrier. STEM CELLS 36, 1816–1827.

DeStefano J. G., Xu Z. S., Williams A. J., Yimam N., Searson P. C. (2017) Effect of shear stress on iPSC-derived human brain microvascular endothelial cells (dhBMECs). Fluids Barriers CNS 14, 20.

Di Marco A., Vignone D., Gonzalez Paz O., Fini I., Battista M. R., Cellucci A., Bracacel E., et al. (2020) Establishment of an in Vitro Human Blood-Brain Barrier Model Derived from Induced Pluripotent Stem Cells and Comparison to a Porcine Cell-Based System. Cells 9, 994.

Helms H. C., Abbott N. J., Burek M., Cecchelli R., Couraud P.-O., Deli M. A., Förster C., et al. (2016) In vitro models of the blood–brain barrier: An overview of commonly used brain endothelial cell culture models and guidelines for their use. J. Cereb. Blood Flow Metab. 36, 862–890.

Hollmann E. K., Bailey A. K., Potharazu A. V., Neely M. D., Bowman A. B., Lippmann E. S. (2017) Accelerated differentiation of human induced pluripotent stem cells to blood–brain barrier endothelial cells. Fluids Barriers CNS 14, 9.

Katt M. E., Linville R. M., Mayo L. N., Xu Z. S., Searson P. C. (2018) Functional brain-specific microvessels from iPSC-derived human brain microvascular endothelial cells: the role of matrix composition on monolayer formation. Fluids Barriers CNS 15, 7.

Katt M. E., Xu Z. S., Gerecht S., Searson P. C. (2016) Human Brain Microvascular Endothelial Cells Derived from the BC1 iPS Cell Line Exhibit a Blood–Brain Barrier Phenotype. PLOS ONE 11, e0152105.

Köster J., Rahmann S. (2018) Snakemake-a scalable bioinformatics workflow engine. Bioinforma. Oxf. Engl. 34, 3600.

Kumar K. K., Lowe, Jr. E. W., Aboud A. A., Neely M. D., Redha R., Bauer J. A., Odak M., et al. (2014) Cellular manganese content is developmentally regulated in human dopaminergic neurons. Sci. Rep. 4, 6801.

Liebner S., Corada M., Bangsow T., Babbage J., Taddei A., Czupalla C. J., Reis M., et al. (2008) Wnt/β-catenin signaling controls development of the blood–brain barrier. J. Cell Biol. 183, 409–417.

Lim R. G., Quan C., Reyes-Ortiz A. M., Lutz S. E., Kedaigle A. J., Gipson T. A., Wu J., et al. (2017) Huntington’s Disease iPSC-Derived Brain Microvascular Endothelial Cells Reveal WNT-Mediated Angiogenic and Blood–Brain Barrier Deficits. Cell Rep. 19, 1365–1377.

Linville R. M., DeStefano J. G., Sklar M. B., Xu Z., Farrell A. M., Bogorad M. I., Chu C., et al. (2019) Human iPSC-derived blood–brain barrier microvessels: validation of barrier function and endothelial cell behavior. Biomaterials 190–191, 24–37.

Lippmann E. S., Al-Ahmad A., Azarin S. M., Palecek S. P., Shusta E. V. (2014a) A retinoic acid-enhanced, multicellular human blood-brain barrier model derived from stem cell sources. Sci. Rep. 4, 4160.

Lippmann E. S., Azarin S. M., Kay J. E., Nessler R. A., Wilson H. K., Al-Ahmad A., Palecek S. P., Shusta E. V. (2012) Derivation of blood-brain barrier endothelial cells from human pluripotent stem cells. Nat. Biotechnol. 30, 783–791.

Lippmann E. S., Azarin S. M., Palecek S. P., Shusta E. V. (2020) Commentary on human pluripotent stem cell-based blood-brain barrier models. Fluids Barriers CNS 17, 64.

Lippmann E. S., Estevez-Silva M. C., Ashton R. S. (2014b) Defined Human Pluripotent Stem Cell Culture Enables Highly Efficient Neuroepithelium Derivation Without Small Molecule Inhibitors. Stem Cells 32, 1032–1042.

Love M. I., Huber W., Anders S. (2014) Moderated estimation of fold change and dispersion for RNA-seq data with DESeq2. Genome Biol. 15, 550.

Mantle J. L., Min L., Lee K. H. (2016) Minimum Transendothelial Electrical Resistance Thresholds for the Study of Small and Large Molecule Drug Transport in a Human in Vitro Blood-Brain Barrier Model. Mol. Pharm. 13, 4191–4198.

Motallebnejad P., Thomas A., Swisher S. L., Azarin S. M. (2019) An isogenic hiPSC-derived BBB-on-a-chip. Biomicrofluidics 13, 064119.

Neal E. H., Marinelli N. A., Shi Y., McClatchey P. M., Balotin K. M., Gullett D. R., Hagerla K. A., et al. (2019) A Simplified, Fully Defined Differentiation Scheme for Producing Blood-Brain Barrier Endothelial Cells from Human iPSCs. Stem Cell Rep. 12, 1380–1388.

Ohshima M., Kamei S., Fushimi H., Mima S., Yamada T., Yamamoto T. (2019) Prediction of Drug Permeability Using In Vitro Blood-Brain Barrier Models with Human Induced Pluripotent Stem Cell-Derived Brain Microvascular Endothelial Cells. BioResearch Open Access 8, 200–209.

Park T.-E., Mustafaoglu N., Herland A., Hasselkus R., Mannix R., FitzGerald E. A., Prantil-Baun R., et al. (2019) Hypoxia-enhanced Blood-Brain Barrier Chip recapitulates human barrier function and shuttling of drugs and antibodies. Nat. Commun. 10, 1–12.

Roux G. L., Jarray R., Guyot A.-C., Pavoni S., Costa N., Théodoro F., Nassor F., et al. (2019) Proof-of-Concept Study of Drug Brain Permeability Between in Vivo Human Brain and an in Vitro iPSCs-Human Blood-Brain Barrier Model. Sci. Rep. 9, 16310.

Schindelin J., Arganda-Carreras I., Frise E., Kaynig V., Longair M., Pietzsch T., Preibisch S., et al. (2012) Fiji: an open-source platform for biological-image analysis. Nat. Methods 9, 676–682.

Stebbins M. J., Wilson H. K., Canfield S. G., Qian T., Palecek S. P., Shusta E. V. (2016) Differentiation and characterization of human pluripotent stem cell-derived brain microvascular endothelial cells. Methods 101, 93–102.

Stenman J. M., Rajagopal J., Carroll T. J., Ishibashi M., McMahon J., McMahon A. P. (2008) Canonical Wnt Signaling Regulates Organ-Specific Assembly and Differentiation of CNS Vasculature. Science 322, 1247–1250.

Tidball A. M., Neely M. D., Chamberlin R., Aboud A. A., Kumar K. K., Han B., Bryan M. R., Aschner M., Ess K. C., Bowman A. B. (2016) Genomic Instability Associated with p53 Knockdown in the Generation of Huntington’s Disease Human Induced Pluripotent Stem Cells. PLOS ONE 11, e0150372.

Vanlandewijck M., He L., Mäe M. A., Andrae J., Ando K., Del Gaudio F., Nahar K., et al. (2018) A molecular atlas of cell types and zonation in the brain vasculature. Nature 554, 475–480.

Vatine G. D., Al-Ahmad A., Barriga B. K., Svendsen S., Salim A., Garcia L., Garcia V. J., et al. (2017) Modeling Psychomotor Retardation using iPSCs from MCT8-Deficient Patients Indicates a Prominent Role for the Blood-Brain Barrier. Cell Stem Cell 20, 831–843.

Vatine G. D., Barrile R., Workman M. J., Sances S., Barriga B. K., Rahnama M., Barthakur S., et al. (2019) Human iPSC-Derived Blood-Brain Barrier Chips Enable Disease Modeling and Personalized Medicine Applications. Cell Stem Cell 24, 995–1005.e6.

Verheyen E. M., Gottardi C. J. (2010) Regulation of Wnt/β-Catenin Signaling by Protein Kinases. Dev. Dyn. Off. Publ. Am. Assoc. Anat. 239, 34–44.

Wang Y. I., Abaci H. E., Shuler M. L. (2016) Microfluidic blood-brain barrier model provides in vivo-like barrier properties for drug permeability screening. Biotechnol. Bioeng. 114, 184–194.

Yousef H., Czupalla C. J., Lee D., Chen M. B., Burke A. N., Zera K. A., Zandstra J., et al. (2019) Aged blood impairs hippocampal neural precursor activity and activates microglia via brain endothelial cell VCAM1. Nat. Med. 25, 988–1000.

Zhang S. L., Yue Z., Arnold D. M., Artiushin G., Sehgal A. (2018) A circadian clock in the blood-brain barrier regulates xenobiotic efflux. Cell 173, 130–139.e10.

