## Supplemental information for "Influence of basal media composition on barrier fidelity within human pluripotent stem cell-derived blood-brain barrier models"

**Table S1. Primary antibodies used for immunocytochemistry.**

| **Target** | **Antibody Species** | **Vendor** | **Clone or product number** | **Dilution** |
| --- | --- | --- | --- | --- |
| Claudin-5 | Mouse | Thermo Fisher Scientific | 4C3C2 | 1:50 |
| Occludin | Mouse | Thermo Fisher Scientific | OC-3F10 | 1:100 |
| VE-cadherin | Goat | R&D Systems | AF938 | 1:100 |
| GLUT-1 | Mouse | Thermo Fisher Scientific | SPM498 | 1:50 |

**Table S2. Secondary antibodies used for immunocytochemistry.**

| **Species Reactivity** | **Host** | **Conjugate** | **Vendor** | **Dilution** |
| --- | --- | --- | --- | --- |
| Mouse | Donkey | Alexa Fluor 555 | Thermo Fisher Scientific | 1:200 |
| Goat | Donkey | Alexa Fluor 488 | Thermo Fisher Scientific | 1:200 |

**Table S3. Primary antibodies used for western blotting.**

| **Target** | **Antibody Species** | **Vendor** | **Clone or product number** | **Dilution** |
| --- | --- | --- | --- | --- |
| FAK | Rabbit | Cell Signaling | 3285 | 1:1000 |
| Phospho-FAK (Tyr397) | Rabbit | Cell Signaling | 3283 | 1:1000 |
| ERK1/2 | Rabbit | R&D Systems | AF1576 | 1:1000 |
| Phospho-ERK1(T202/Y204)/  ERK2(T185/Y187) | Rabbit | R&D Systems | AF1018 | 1:2000 |

**Table S4. Full list of differentially expressed genes in DMEM versus NB basal media.**

This table is attached as an Excel spreadsheet.

**Table S5. Full list of altered KEGG pathways in DMEM versus NB basal media.**

| **Description** | **Change** | **GeneRatio** | **qvalue** |
| --- | --- | --- | --- |
| Axon guidance | Upregulated | 52/1018 | 1.35E-06 |
| Focal adhesion | Upregulated | 53/1018 | 9.61E-06 |
| Basal cell carcinoma | Upregulated | 23/1018 | 9.23E-05 |
| Human papillomavirus infection | Upregulated | 71/1018 | 2.00E-04 |
| Wnt signaling pathway | Upregulated | 41/1018 | 2.57E-04 |
| ECM-receptor interaction | Upregulated | 27/1018 | 2.67E-04 |
| Melanogenesis | Upregulated | 29/1018 | 4.44E-04 |
| Hepatocellular carcinoma | Upregulated | 41/1018 | 5.81E-04 |
| Protein digestion and absorption | Upregulated | 28/1018 | 1.39E-03 |
| Breast cancer | Upregulated | 36/1018 | 1.39E-03 |
| Gastric cancer | Upregulated | 36/1018 | 1.71E-03 |
| mTOR signaling pathway | Upregulated | 36/1018 | 3.70E-03 |
| ErbB signaling pathway | Upregulated | 23/1018 | 5.06E-03 |
| Cushing syndrome | Upregulated | 35/1018 | 6.92E-03 |
| Hippo signaling pathway | Upregulated | 35/1018 | 8.33E-03 |
| Lysosome | Upregulated | 29/1018 | 1.78E-02 |
| Signaling pathways regulating pluripotency of stem cells | Upregulated | 31/1018 | 2.47E-02 |
| GnRH secretion | Upregulated | 17/1018 | 2.78E-02 |
| Steroid biosynthesis | Upregulated | 8/1018 | 2.78E-02 |
| Endocrine resistance | Upregulated | 23/1018 | 2.79E-02 |
| Rap1 signaling pathway | Upregulated | 41/1018 | 3.30E-02 |
| N-Glycan biosynthesis | Upregulated | 14/1018 | 3.38E-02 |
| Proteoglycans in cancer | Upregulated | 40/1018 | 3.42E-02 |
| EGFR tyrosine kinase inhibitor resistance | Upregulated | 19/1018 | 3.98E-02 |
| Calcium signaling pathway | Upregulated | 45/1018 | 3.98E-02 |
| Phosphatidylinositol signaling system | Upregulated | 22/1018 | 4.16E-02 |
| Lysine degradation | Upregulated | 16/1018 | 4.18E-02 |
| Glioma | Upregulated | 18/1018 | 4.48E-02 |
| Notch signaling pathway | Upregulated | 14/1018 | 4.48E-02 |
| Pathways of neurodegeneration - multiple diseases | Upregulated | 79/1018 | 4.48E-02 |
| Regulation of actin cytoskeleton | Upregulated | 41/1018 | 4.48E-02 |
| Mannose type O-glycan biosynthesis | Upregulated | 8/1018 | 4.54E-02 |
| Cell cycle | Upregulated | 26/1018 | 4.69E-02 |
| Glycosaminoglycan degradation | Upregulated | 7/1018 | 5.08E-02 |
| Hedgehog signaling pathway | Upregulated | 13/1018 | 5.90E-02 |
| Prostate cancer | Upregulated | 21/1018 | 6.39E-02 |
| Glycosaminoglycan biosynthesis - chondroitin sulfate / dermatan sulfate | Upregulated | 7/1018 | 6.42E-02 |
| Alzheimer disease | Upregulated | 62/1018 | 7.05E-02 |
| Cell adhesion molecules | Upregulated | 29/1018 | 7.41E-02 |
| Herpes simplex virus 1 infection | Downregulated | 145/1047 | 1.12E-20 |
| p53 signaling pathway | Downregulated | 22/1047 | 1.09E-02 |
| Autophagy - animal | Downregulated | 34/1047 | 1.09E-02 |
| Platinum drug resistance | Downregulated | 21/1047 | 2.00E-02 |
| Basal transcription factors | Downregulated | 15/1047 | 2.12E-02 |
| Peroxisome | Downregulated | 21/1047 | 8.46E-02 |
| SNARE interactions in vesicular transport | Downregulated | 11/1047 | 9.28E-02 |

**Figure S1. Raw images of the phospho-kinase array and array coordinates.**

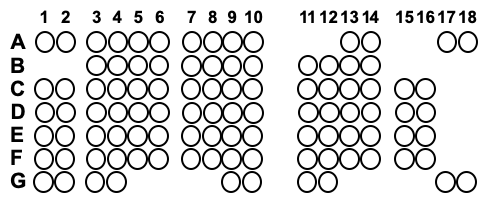

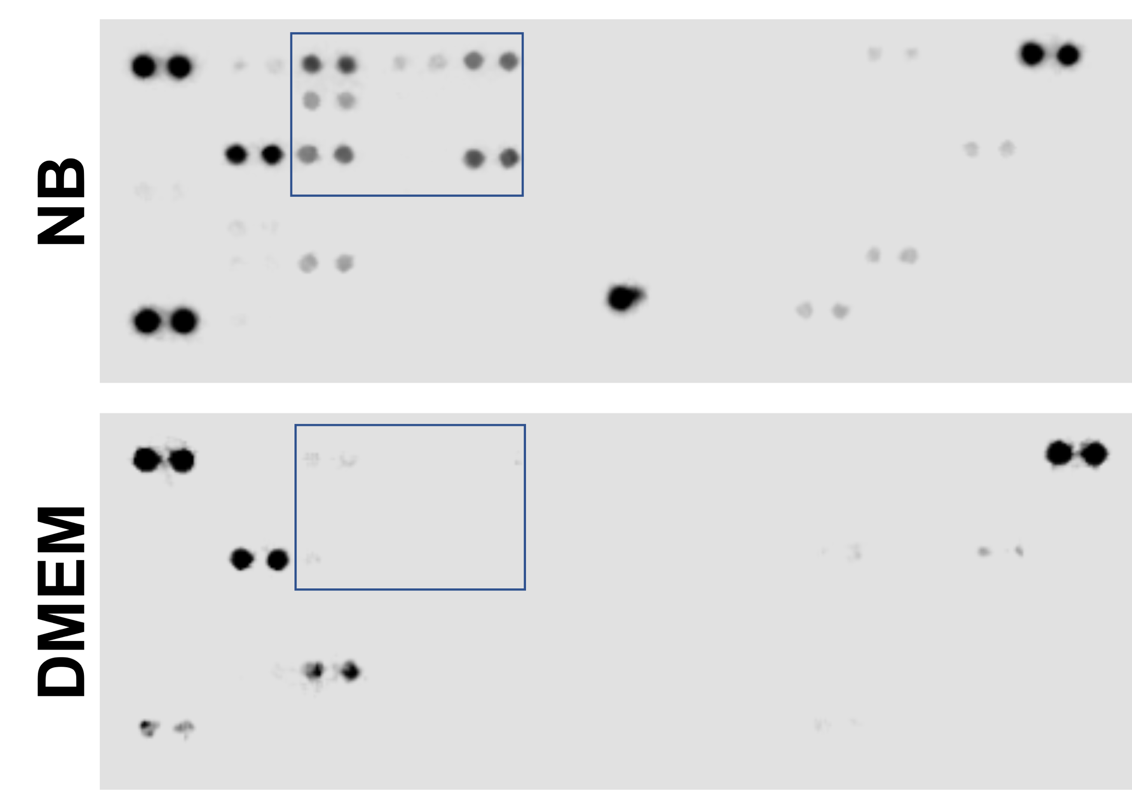

| **Coordinate** | **Target** | **Phosphorylation site** | **Coordinate** | **Target** | **Phosphorylation site** |
| --- | --- | --- | --- | --- | --- |
| A1, A2 | Reference spot | −−− | D9, D10 | STAT5a | Y694 |
| A3, A4 | p38⍺ | T180/Y182 | D11, D12 | p70 S6 kinase | T421/S424 |
| A5, A6 | ERK1/2 | T202/Y204, T185/Y187 | D13, D14 | RSK1/2/3 | S380/S386/S377 |
| A7, A8 | JNK1/2/3 | T183/Y185, T221/Y223 | D15, D16 | eNOS | S1177 |
| A9, A10 | GSK-3⍺/β | S21/S9 | E1, E2 | Fyn | Y420 |
| A13, A14 | p53 | S392 | E3, E4 | Yes | Y426 |
| A17, A18 | Reference spot | −−− | E5, E6 | Fgr | Y412 |
| B3, B4 | EGFR | Y1086 | E7, E8 | STAT6 | Y641 |
| B5, B6 | MSK1/2 | S376/S360 | E9, E10 | STAT5b | Y699 |
| B7, B8 | AMPK⍺1 | T183 | E11, E12 | STAT3 | Y705 |
| B9, B10 | Akt1/2/3 | S473 | E13, E14 | p27 | T198 |
| B11, B12 | Akt1/2/3 | T308 | E15, E16 | PLC-ɣ1 | Y783 |
| B13, B14 | p53 | S46 | F1, F2 | Hck | Y411 |
| C1, C2 | TOR | S2448 | F3, F4 | Chk-2 | T68 |
| C3, C4 | CREB | S133 | F5, F6 | FAK | Y397 |
| C5, C6 | HSP27 | S78/S82 | F7, F8 | PDGFRβ | Y751 |
| C7, C8 | AMPK⍺2 | T172 | F9, F10 | STAT5a/b | Y694/Y699 |
| C9, C10 | β-catenin | −−− | F11, F12 | STAT3 | S727 |
| C11, C12 | p70 S6 kinase | T389 | F13, F14 | WNK1 | T60 |
| C13, C14 | p53 | S15 | F15, F16 | PYK2 | Y402 |
| C15, C16 | c-Jun | S63 | G1, G2 | Reference spot | −−− |
| D1, D2 | Src | Y419 | G3, G4 | PRAS40 | T246 |
| D3, D4 | Lyn | Y397 | G9, G10 | PBS (neg control) | −−− |
| D5, D6 | Lck | Y394 | G11, G12 | HSP60 | −−− |
| D7, D8 | STAT2 | Y689 | G17, G18 | PBS (neg control) | −−− |

The blue box represents the location of the blot that was presented in the main text of this manuscript. The array coordinates and targets were provided on the manufacturer’s website and are reproduced here for convenience.

**Figure S2. Raw images of western blots.**

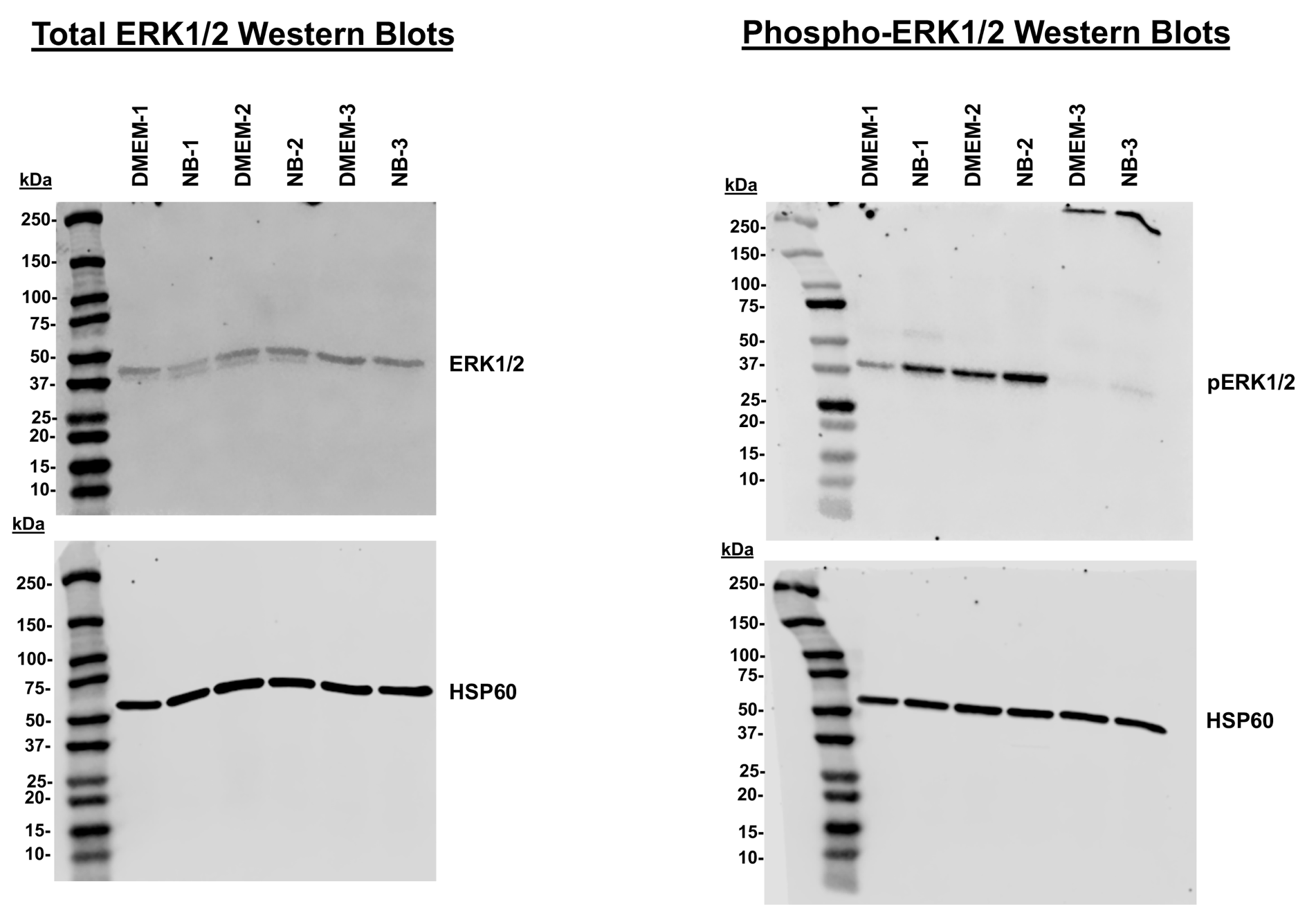

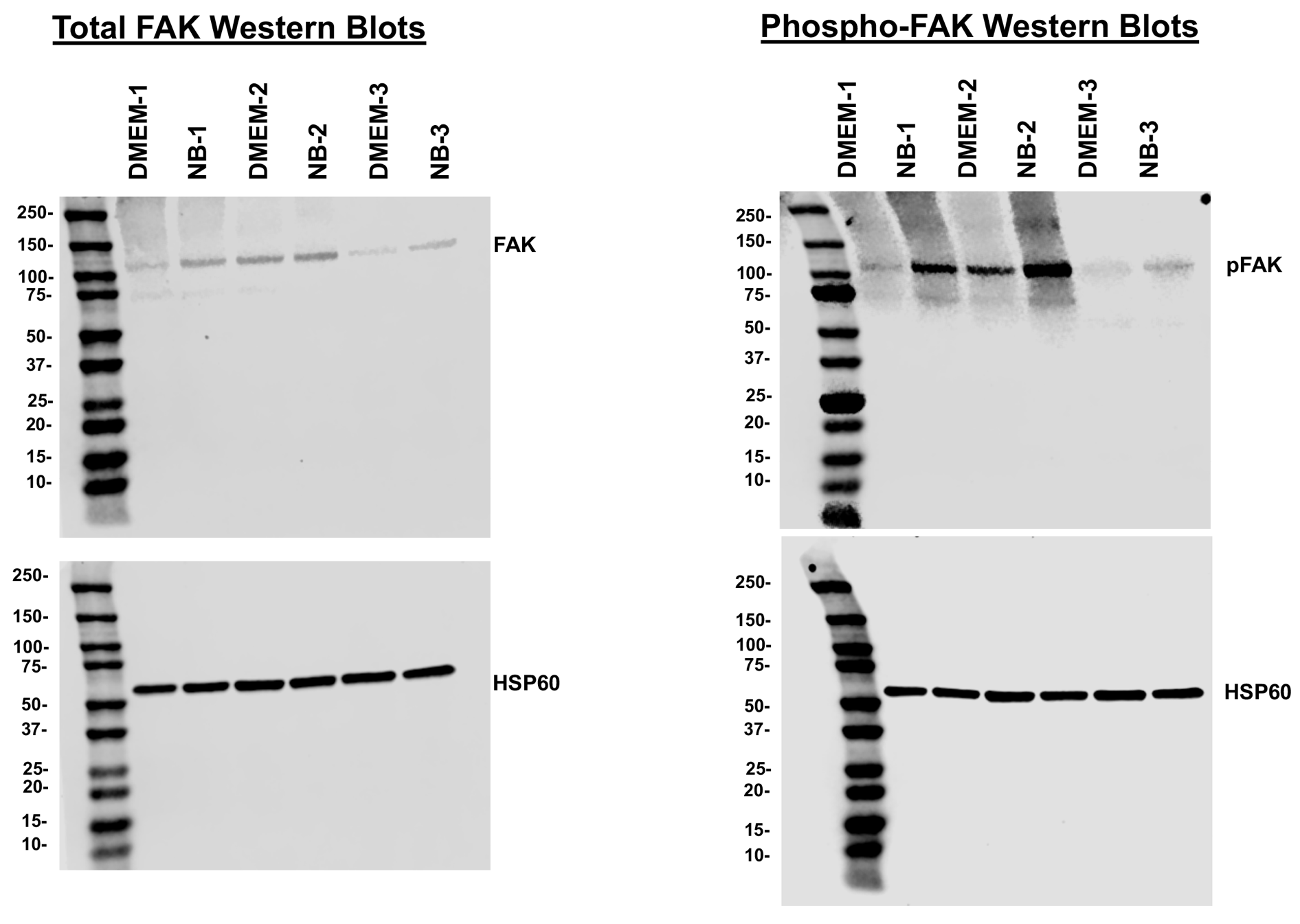
